# Identification and characterization of emNagII, a novel β-1,2-N-acetylglucosaminidase from *Elizabethkingia meningoseptica*

**DOI:** 10.1101/2025.10.15.682710

**Authors:** Shaoxian Lyu, Yongliang Tong, Qiange Lin, Yilin Ye, Xinliu Geng, Cuiying Chen, Xinrong Lu, Guiqin Sun, Li Chen

## Abstract

N-Acetylglucosamine exists in various forms and linkage patterns within organisms, playing a crucial role in numerous vital biological processes. However, research focusing on β-1,2-linked N-acetylglucosamine within complex biantennary N-glycans on the cell surface remain largely unexplored. Currently, there is a lack of efficient tools and methods capable of directly modifying terminal N-acetylglucosamine on living cells. In this study, we identified a novel β-N-acetylhexosaminidase from *Elizabethkingia meningoseptica*, which demonstrated favorable enzymatic stability within a pH range of 5-8 and at temperatures from 4 °C to 37 °C. This enzyme specifically cleaves non-reducing terminal β-1,2-N-acetylglucosamine, enabling direct removal of this modification from oligosaccharides and native glycoproteins. More importantly, we demonstrate for the first time that this enzyme successfully removes β-1,2-linked N-acetylglucosamine on living cell surfaces. Given its microbial origin and potential utility in living cell glycan editing, we have named it emNagII or cell-surface glycan-editing N-acetylglucosaminidase (csgeNagII). Through sequence analysis and alanine scanning mutagenesis, we identified a predicted active pocket containing the catalytic residue pair Asp317-Glu318, with Asp317 being essential for enzymatic activity. In summary, our findings provide an effective and reliable method for the targeted removal of β-1,2-N-acetylglucosamine on living cell surfaces, establishing a foundation for further functional studies and practical applications.

## 1. Introduction

N-Acetylglucosamine (GlcNAc) is a common carbohydrate which is formed by a 2-acetylamino-2-deoxyderivative of glucose. It serving as a nutrient source [1], participating in signal transduction [2], immune regulation and disease progression [3-5]. N-Glycosylation is a critical post-translational modification in most eukaryotic cells and essential for protein stability, functionality, and intercellular communication [6-8]. GlcNAc constitutes a core component of N-linked core pentasaccharide and additionally branches into diverse N-glycans structure via β-1,2-, β-1,4-, and β-1,6-linkages [9]. Nevertheless, the functional significance of GlcNAc within N-glycans remains uncharacterized, particularly for β-1,2-linked N-acetylglucosamine (β-1,2-GlcNAc) in complex-type N-glycans. Current limitations in efficient enzymatic tools further impede the decoding of glycan structure information.

β-N-Acetylglucosaminidase is an enzyme that catalyzes the removal of terminal β-linked N-acetylglucosamine (GlcNAc) residues. It belongs to the β-N-acetylhexosaminidase family (EC 3.2.1.52). Systematic research on these enzymes began in 1970, when Matsushima and Yamamoto first isolated a β-N-acetylglucosaminidase from Takadiastase and performed its detailed biochemical characterization [10, 11]. Based on protein sequence similarities, β-N-acetylhexosaminidases are classified into glycoside hydrolase families GH3, GH20, GH84 and GH116 [12-16]. Among these, the GH20 family exhibits the highest structural and functional diversity of β-N-acetylhexosaminidases and represents the extensively characterized subgroup [17, 18].

The β-N-acetylhexosaminidase is commonly employed in the glycan industry to produce GlcNAc from glycan substrates. However, there are relatively few reports on its direct action on native glycoproteins. In 2018, Bulter et al. reported that β-N-acetylhexosaminidase from *Streptococcus pneumoniae* can cleave terminal GlcNAc of the native N-glycan structure of antibodies, enabling the modification of antibody glycoforms [19, 20]. Although antibodies harbor significant proportion of G0 and G0F glycoforms [21], the biological significance of terminal β-1,2-GlcNAc residues remains poorly. Cell surfaces are densely populated with glycoconjugates, some of which also display terminal β-1,2-GlcNAc moieties [22, 23]. Nevertheless, research on the function of cell surface terminal β-1, 2-GlcNAc remains limited, partly due to the lack of efficient tools to cleave GlcNAc on living cells.

In this study, we isolated a β-N-acetylhexosaminidase from the genome of *Elizabethkingia meningoseptica* and verified its enzymatic activity toward terminal β-1,2-GlcNAc on oligosaccharides, glycoproteins, and cell surface glycans. It was named emNagII (csgeNagII) as a novel tool for decoding the specific roles and functions of β-1,2-GlcNAc signals in the natural antibodies and on living cell surfaces.

## 2. Materials and methods

### 2.1. Bioinformatics analysis of emNagII – a putative β-N-acetylhexosaminidase from E. meningoseptica

The gene encoding emNagII was obtained from the *E. meningoseptica* FMS-007, a clinically isolated strain from a T-cell non-Hodgkin’s lymphoma patient. Reference β-N-acetylhexosaminidase sequences were retrieved from the CAZy database and Uniprot for phylogenetic analysis and multiple-sequence alignment. A phylogenetic tree was built with MEGA (version 12) using the neighbor-joining method. Multiple sequence alignment was performed with ClustalW and ESPript (version 3.0). The signal peptide was predicted using SignalP (version 5.0). Structural features of emNagII were analyzed via the InterPro database (https://www.ebi.ac.uk/interpro/), and its three-dimensional was modeled with AlphaFold2 and displayed by PyMOL. Molecular docking was conducted by MOE software.

### 2.2. Cloning and protein expression

The *emNagII* gene without the signal peptide was amplified by PCR from the genomic DNA of strain FMS-007 using primers containing NheI and XhoI restriction sites. The PCR products was ligated into pET28a vector using T4 DNA Ligase (Thermo, #EL0012). All mutants were generated from the wild-type pET28a-emNagII plasmid using the QuickMutation™ Site-Directed Mutagenesis Kit (Beyotime Biotechnology, #D0206S). Both wild-type and mutant plasmids were transformed into *E. coli* BL21 (DE3) (Yeasen, #11804) for protein expression. Single colonies were cultured in Luria-Bertani medium supplemented with 50 μg/mL kanamycin, and protein expression was induced by IPTG. Finally, proteins were purified using Ni Sepharose™ 6 Fast Flow (Cytiva, #175318). Endotoxins were removed with a high-capacity endotoxin removal spin kit (Thermo Fisher, #88275).

### 2.3. Enzymatic assay against p-nitrophenyl glycosides

To determine substrate specificity, emNagII was incubated with various p-nitrophenyl (pNP) α/β-glycosides at 37 ℃ for 1 h, and reactions were terminated by adding Na_2_CO_3_. Optimal pH was assessed by measuring activity in citrate (50 mM, pH 3.0-7.0) or Tris-HCl (50 mM, pH 8.0-10.0) buffer after 5 min at 37 ℃. pH stability was evaluated by pre-incubating emNagII in buffers of different pH for 1 h before adding substrate at the optimal pH. For determine the optimal temperature of emNagII, setting the reaction systems at 4, 20, 25, 30, 37, 45, 50, 55, 65 and 75 ℃, separately. All reactions were terminated after incubating 5 min. The temperature tolerance was assessed by pre-incubating emNagII at each temperature for 1 h prior to activity measurement at optimal temperature. The effects of metal ions (Zn^2+^, Ca^2+^, Ni^2+^, Cu^2+^, Mn^2+^, Mg^2+^ and Fe^2+^) and chemical reagents (UREA, SDS and EDTA) were tested by adding each compound to the reaction mixture. Absorbance was measured at 405 nm after termination. All experiments were performed in triplicated.

### 2.4. Enzyme activity against glycoproteins

IgG with complex N-glycans was selected as the substrate to assess enzymatic activity. A mixture containing 100 μg IgG and emNagII in 200 μL reaction was incubated at 37 ℃ for 12 h, denatured at 100 ℃ for 10 min. Subsequently, 5 μL PNGase F was added to release N-glycans from IgG. The glycans were purified by 3 kDa ultrafiltration, and desalted with a Supelclean™ ENVI-Carb™ SPE cartridge. After vacuum dried, the sample was analyzed by Matrix-Assisted Laser Desorption/Ionization Time-of-Flight (MALDI-TOF) mass spectrometry.

### 2.5. Enzyme activity on living immune cell surfaces

Splenic cells were isolated from adult C57BL/6 mice to examine the enzymatic activity of emNagII on living cell surfaces. All animal procedures were approved by the Ethics Committee of Experimental Research, Fudan University Shanghai Medical College. Single-cell suspensions were prepared by grinding spleens through a 200-mesh strainer. After centrifugation, erythrocytes were lysed using erythrocyte lysis buffer (Yeasen, #40401) for 10 min at room temperature. The remaining cells were washed and resuspended in RPMI-1640 medium at a density of 5×10^6^ cells/mL and divided into groups for enzymatic treatments. Following 4 h incubation, enzymes were removed by centrifugation. Cell surface N-acetylglucosamines were labeled with Wheat germ agglutinin (Vector Laboratories, # B-1025-5) and detected with Alexa Fluor®488 Conjugated anti-His Antibody (Cell Signaling Technology, #14930). Finally, the samples were analyzed by flow cytometry (BD, FACSCalibur).

## 3. Results

### 3.1. Bioinformatics analysis of a novel candidate β-N-Acetylhexosaminidase gene in the whole-genome sequence of E. meningoseptica

The clinical strain of *E. meningoseptica* FMS-007 was isolated from a T-cell non-Hodgkin’s lymphoma patient in our laboratory [24]. Whole-genome bioinformatic analysis identified a β-N-Acetylhexosaminidase candidate named emNagII. This protein consists of 748 amino acids including a signal peptide of 21 amino acids. Phylogenetic analysis of emNagII with other β-N-Acetylhexosaminidase in GeneBank^TM^ indicated that emNagII belonged to the glycoside hydrolase family 20 (GH20). Its closest homolog was the β-N-acetylhexosaminidase (BT0459) from *Bacteroides thetaiotaomicron* (Fig.1), a common gut symbiotic in humans and mice [25]. Similar to *E. meningoseptica*, this bacterium expresses diverse glycosidases, which may be related to its ability to digest dietary polysaccharides and host glycans [26-28].

**Figure 1.**
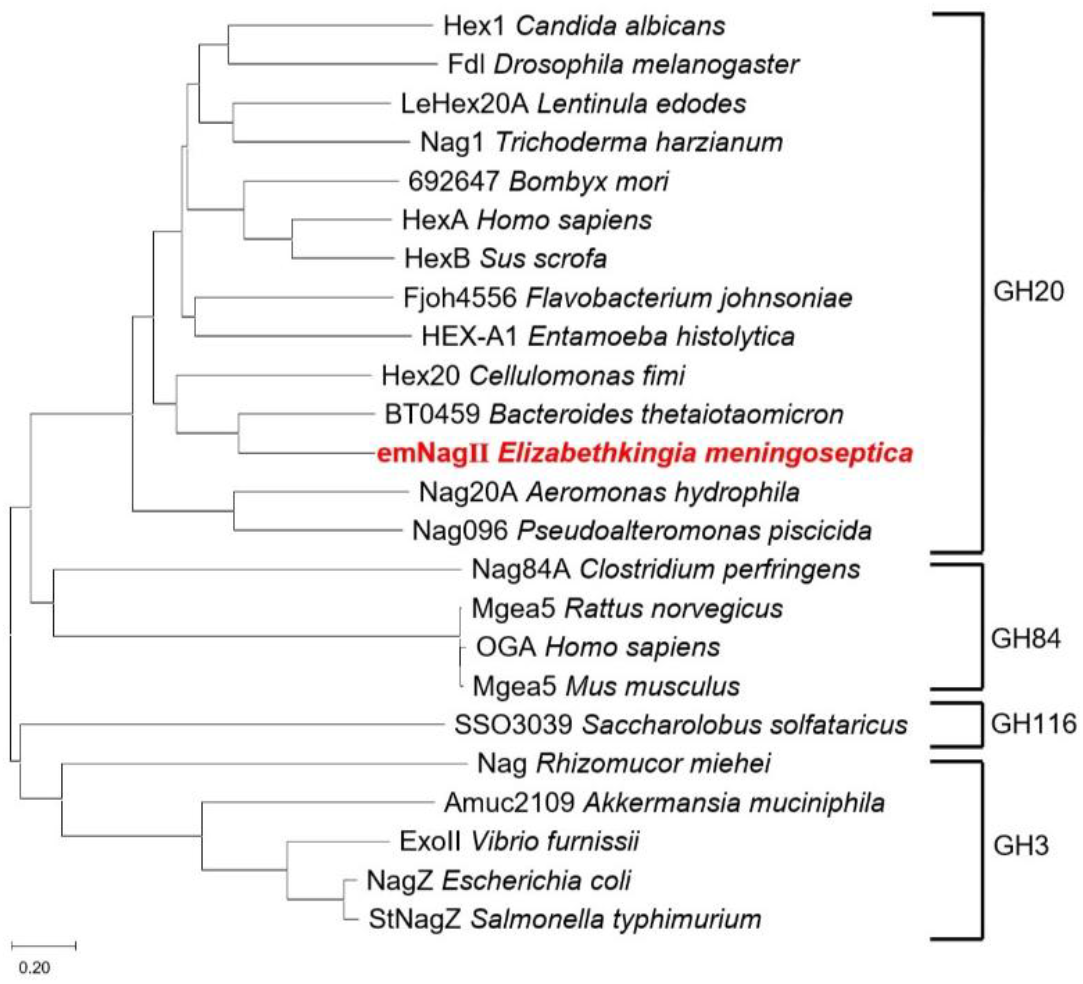
Phylogenetic tree of emNagII and characterized β-N-Acetylhexosaminidase. Amino acid sequences were retrieved from Uniprot database. The phylogenetic tree was constructed using the neighbor joining method. The candidate gene emNagII was highlighted in red font, which was predicted to belong to the GH20 family.

### 3.2. Substrate specificity and β-N-acetylglucosaminidase activity of emNagII on pNPs

The *emNagII* gene excluding the signal peptide was amplified from *E. meningoseptica* FMS-007 genomic DNA and cloned into pET-28a expression vector to produce a N-terminal and C-terminal fusion protein. Purified protein migrated as a single 83.5 kDa band on SDS-PAGE matching the predicted molecular weight (Fig.2a).

**Figure 2.**
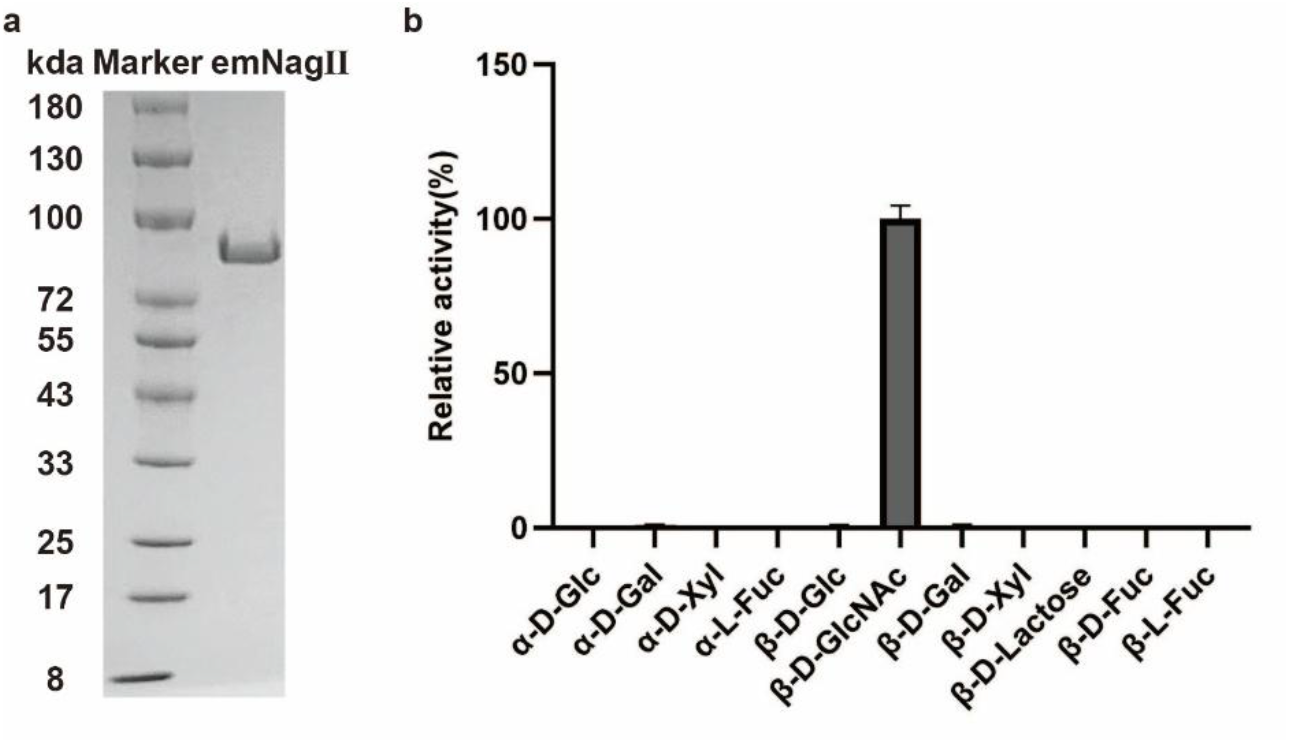
Characterization of emNagII. (a) Coomassie brilliant blue staining verification of purified emNagII. (b) The substrate specificity was verified using 11 p-nitrophenyl α/β-glycosides.

For substrate specificity assay, the enzymatic activity of emNagII against 11 p-nitrophenyl α/β-glycosides was tested. It hydrolyzed pNP-β-D-GlcNAc confirming its function as a β-N-acetylglucosaminidase (Fig.2b). The optimal pH of emNagII was tested to be 5-8 (Fig.3a). Activity declined sharply below pH 4 and was nearly absent at pH 10 (Fig.3b). The optimum temperature of emNagII was 37 ℃ while its activity lost completely above 45 ℃ (Fig.3c-d).

**Figure 3.**
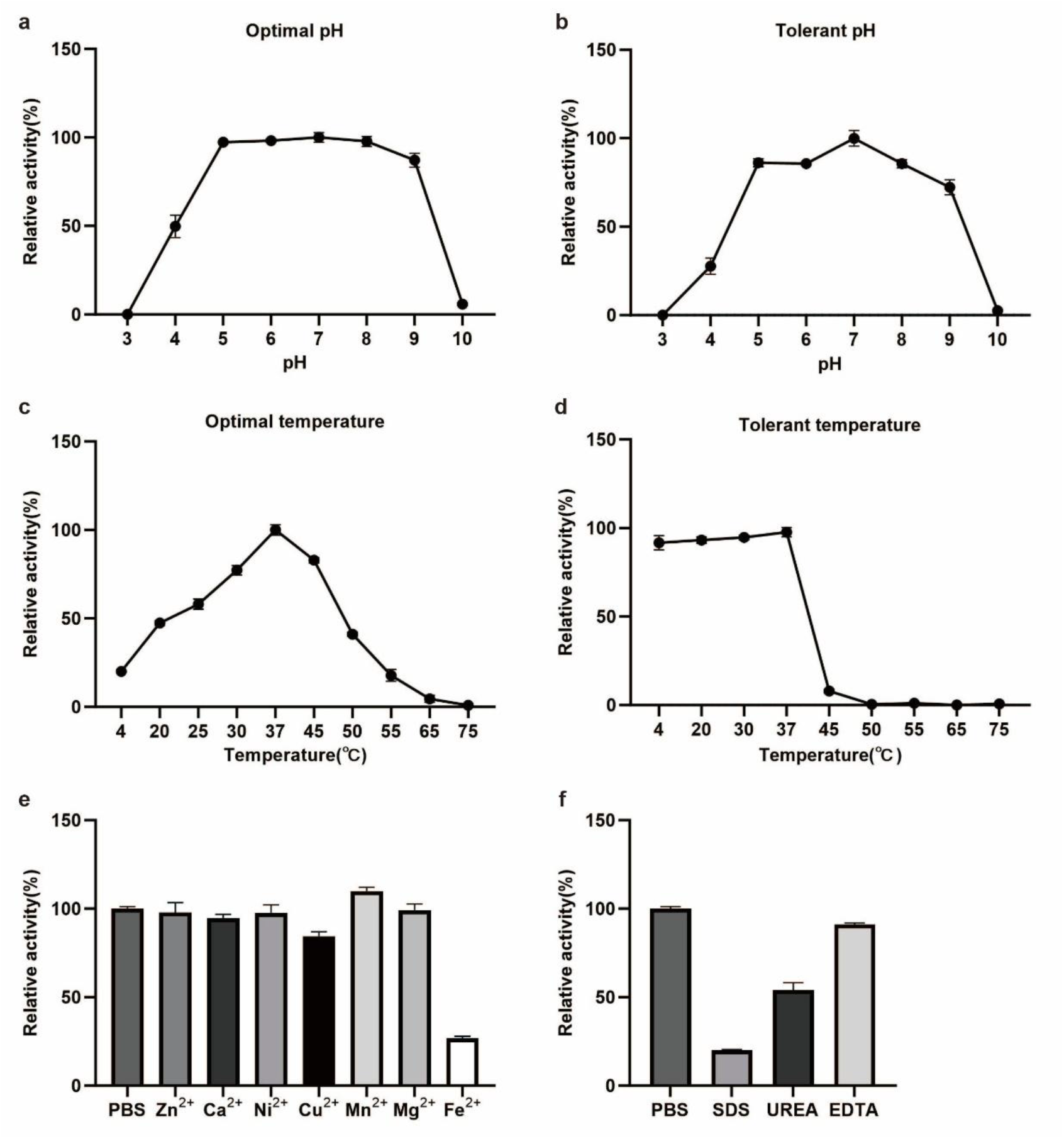
Enzymatic characterization of emNagII. (a) Optimal pH of emNagII. (b) Tolerant pH of emNagII. (c) Optimal temperature of emNagII. (d) Tolerant temperature of emNagII. (e) Effect of metal ions on emNagII activity. (f) Effect of compounds on emNagII activity.

The effects of metal ions were examined at a final concentration of 1 mM. Fe^2+^ strongly inhibited emNagII activity and Cu^2+^ caused mild inhibition. Zn^2+^, Ca^2+^, Ni^2+^, Mn^2+^ and Mg^2+^ showed no effect (Fig.3e). The chelator EDTA at a final concentration of 5 mM had no significant influence on the enzyme activity, whereas 1 M UREA and 0.1 % SDS severely inactivated the enzyme (Fig.3f).

### 3.3. Sequence analysis and defining the enzymatic active sites for β-N-acetylglucosamines hydrolysis of emNagII

Six β-N-acetylglucosaminidases (Hex20, BT0459, HEX-A1, Fjoh4556, HexA, and HexB) from the GH20 family were selected for multiple sequence alignment with emNagII (Fig. S1). The alignment revealed that these enzymes are highly conserved. Residues Arg155, Asp184, His255, Asp317, Glu318, Tyr409, Asp411, Tyr412, and Glu461 were completely or highly conserved among bacterial homologs.

Analysis through the InterPro database indicated that emNagII shares structural features with most GH20 family β-N-acetylhexosaminidases. The structure of emNagII was predicted using the AlphaFold2 model. Its structure could be divided into three parts—a N-terminal domain, a catalytic domain and a C-terminal domain (Fig. S2). Molecular docking of N-acetylglucosamine into the emNagII structure suggested that key residues are spatially clustered, potentially forming an active pocket (Fig. 4a).

**Figure 4.**
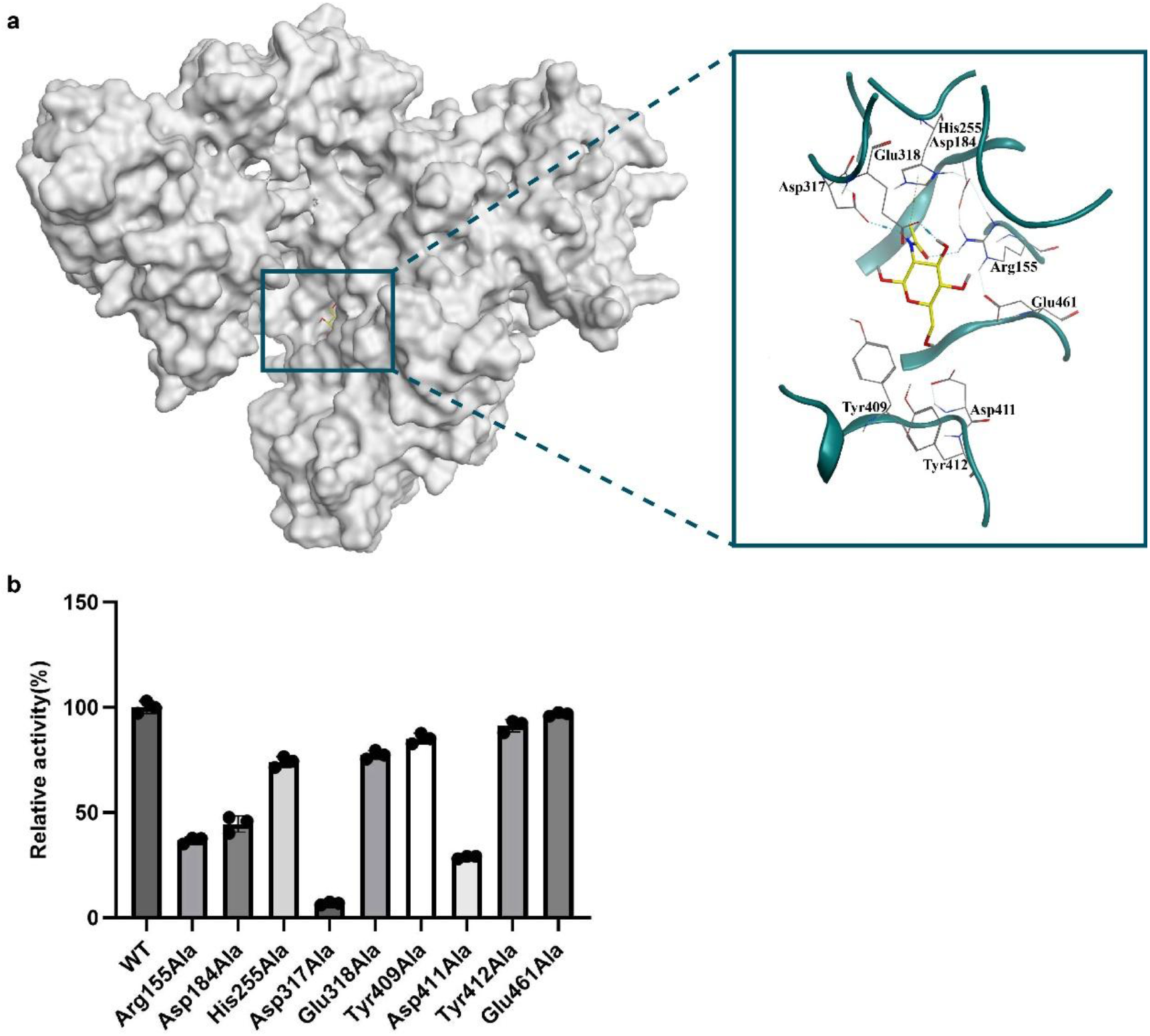
The enzymatic active sites of emNagII. (a) Molecular docking of N-acetylglucosamine with emNagII was predicted using the MOE software. The left picture is the overall prediction result and the right side is the local zoomed-in image. (b) Enzymatic activity against pNP-β-D-GlcNAc of 9 mutants of emNagII were tested.

Based on the result of sequence and structural analyses, Asp184, His255, Asp317, and Glu318 were predicted to be catalytic sites, while Arg155, Tyr409, Asp411, Tyr412, and Glu461 were identified as putative substrate-binding sites. Consistent with the classical mechanism in GH20 enzymes, the conserved Asp-Glu pair (Asp317 and Glu318 in emNagII) is critical for catalysis. Glu318 likely acts as the catalytic acid/base, and Asp317 likely stabilizes the oxazoline intermediate [29, 30].

A series of emNagII mutants were constructed and tested for enzymatic activity using pNP-β-D-GlcNAc as the substrate. Results showed that mutations at Arg155, Asp184, Asp317, and Asp411 led to a significant or complete loss of activity. In contrast, mutations at His255, Glu318, Tyr409, Tyr412, and Glu461 caused slight or no activity loss (Fig. 4b).

### 3.4. β-N-Acetylglucosaminidase activity of emNagII on glycoprotein IgG

To identify the bond type specificity, emNagII was test on various oligosaccharides with different GlcNAc linkages. Results of capillary electrophoresis showed that emNagII specifically hydrolyzes β-1,2-glycosidic bonds (data not shown).

IgG is produced using CHO cell line and contains three major complex N-glycan glycoforms, including G0F, G1F and G2F glycoforms [31]. MALDI-TOF mass spectrometry verified the cleavage of terminal β-1,2-GlcNAcs on IgG by emNagII (Fig.5). After treatment with 100 μg/mL emNagII for 12 h, the ion peaks of G0F (m/z 1485.480) and G1F (m/z 1647.532) glycoforms decreased significantly. Concurrently, new peaks emerged at m/z 1282.419 and m/z 1444.479, consistent with the expected enzymatic digestion products G0F-N and G1F-N glycoforms. Increasing emNagII concentration to 500 μg/mL enhanced substrate reduction and nearly doubled product yield. These results confirming emNagII effectively removes terminal β-1,2-GlcNAcs from native antibodies.

**Figure 5.**
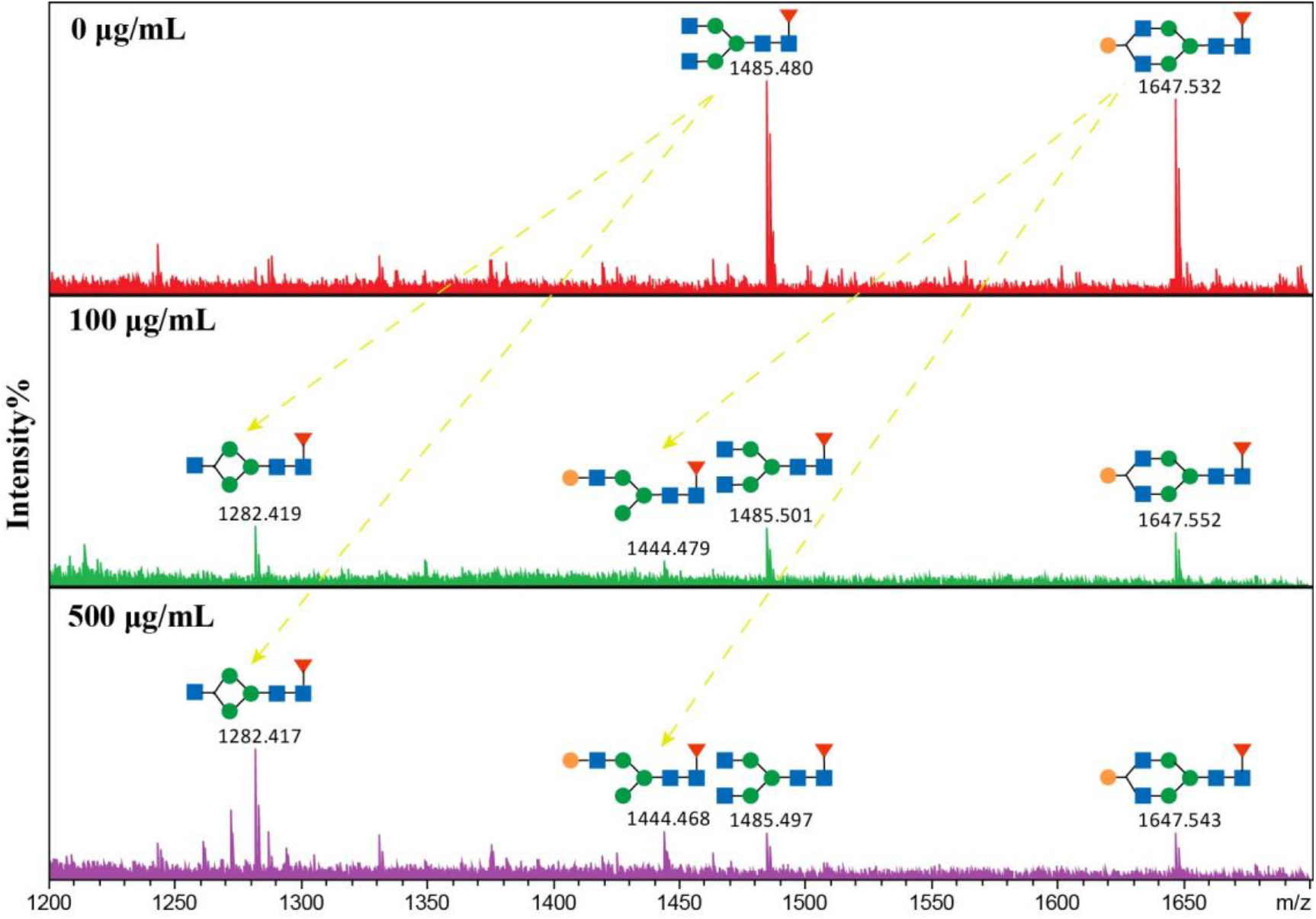
Releasing of the terminal β-N-acetylglucosamine of IgG by emNagII. IgG was incubated with serial concentrations of emNagII. And N-glycan were subsequently collected by PNGase F treatment and detected by MALDI-TOF mass spectrometry. All peaks were annotated with GlycoMod server, and molecular ions were present in sodiated form ([M + Na]^+^).

### 3.5. Capacity of emNagII to cleave terminal β-1,2-GlcNAc on cell surfaces

Previous studies have shown that immune cell surfaces feature complex N-glycans terminating with α-sialic acid, β-1,4-galactose and β-1,2-GlcNAc. To evaluate emNagII activity on living cell surfaces, SiaT and emGalaseE were used to remove sialic acid and galactose first to exposure β-1,2-GlcNAcs (Fig.6a). Wheat Germ Agglutinin (WGA) are reported specifically recognized N-acetylglucosamines [32]. Flow cytometry results showed an increase in the mean fluorescence intensity (MFI) of WGA after treatment with SiaT and emGalaseE [33, 34], indicating successful exposure of β-1,2-GlcNAcs. When added emNagII to co-incubated, the MFI of WGA had reduced to the original level. It suggested that emNagII effectively eliminated terminal β-1,2-GlcNAc on living cell surfaces (Fig.6b).

**Figure 6.**
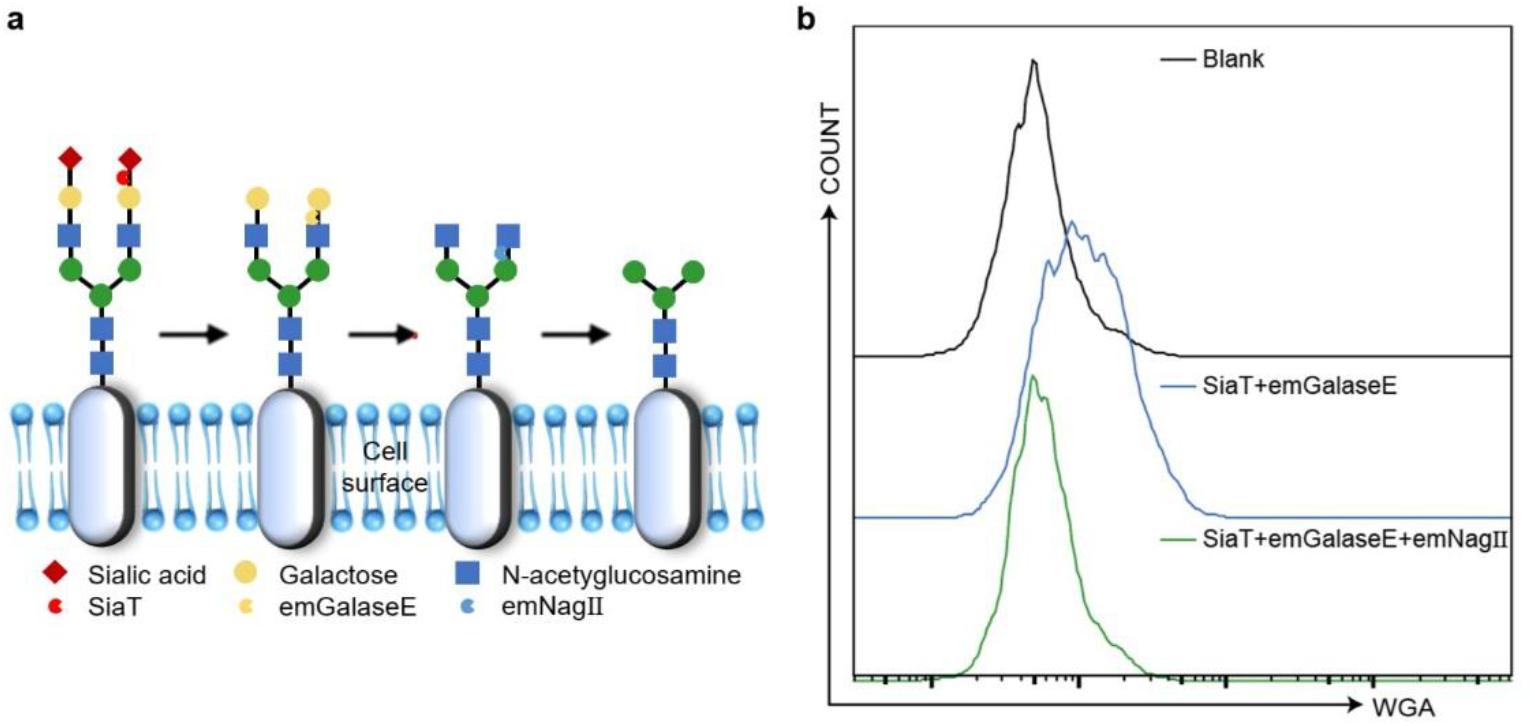
Enzymatic activity of emNagII on living cell surfaces. (a) Schematic diagram of complex N-glycan reconstruction on living cell surfaces N-glycan signals by SiaT, emGalaseE and emNagII. (b) Flow cytometry analysis showed changes in WGA binding fluorescence intensity under different enzymatic treatments on the cell surface.

## 4. Discussion

N-Glycans are ubiquitously distributed on eukaryotic cell surfaces, functioning as crucial molecular interfaces and physical barriers. Besides, these glycans regulate interactions of the coated cells with external factors. The physiological roles of individual monosaccharide constituents within N-glycans require continued mechanistic investigation. Elevated fucosylation and sialylation promote tumorigenesis and metastatic progression by modulating signal transduction, invasive capacity, and angiogenic activity [35-37]. Terminal galactosylated modifies glycans could be recognized by galectins, which participating in the regulation of cell adhesion, phagocytosis, and apoptosis [38-41]. High-mannose N-glycans have also been found to promote tumor stem cell self-renewal and immune evasion by binding to mannose receptors [42]. However, the biological role of GlcNAc on cell surface remains to be illustrated.

The signaling complexity of GlcNAc within N-glycan arises from its occupancy of multiple glycosylation sites and diverse glycosidic linkages (e.g., β-1,2-, β-1,4-, and β-1,6-GlcNAc). On cell surface glycoproteins, the β-1,6-GlcNAc branch facilitates multivalent interactions with galectins and enhances glycoprotein conformational stability [43]. Elevated levels of β-1,6-GlcNAc branch in tumor tissues are associated with metastasis and poor prognosis, whereas β-1,4-linked bisecting GlcNAc counteracts this effect by inhibiting the formation of β-1,6-GlcNAc branch [44]. Bisecting GlcNAc is a specialized N-glycosylation modification which could regulate cell adhesion, fertilization, embryonic development, neurogenesis, tumor metastasis and progression. It is highly expressed in the brain [45], particularly in neurons, where it performs physiological functions and is pathologically related to Alzheimer’s disease [46]. Additionally, bisecting GlcNAc plays a significant role in immune responses. The content of bisecting GlcNAc on the surface affects macrophage polarization, opening new strategies for colorectal cancer treatment [47]. On IgG antibodies, the terminal β-1,2-GlcNAc N-glycans structure (G0/G0F) has been correlated in progeroid syndromes and various autoimmune/inflammatory diseases. Age-related accumulation of IgG-G0 can lead to inflammation [48, 49]. Nevertheless, research on cell surface-localized β-1,2-GlcNAc modified N-glycans remains limited, primarily due to the lack of effective research methods and practical tools.

Since the first systematic study of Taka-N-acetyl-β-D-glucosaminidase by Matsushima and Yamamoto in the 1970s [10, 11], over 100 enzymes have been predicted or identified as Hex/Nag in the CAZy database. Although some of these enzymes are widely used in the engineering of polysaccharides and antibodies, the applications have so far been confined to cell-free systems. In this study, we isolated a novel β-N-acetylglucosaminidase from *E. meningoseptica* and demonstrated its ability to cleave terminal β-1,2-linked N-acetylglucosamine on oligosaccharide chains, glycoproteins, and cell surfaces. Thus, we named it as emNagII for its isolation and glycan editing function, or cell surface glycan editing β-N-acetylglucosaminidase II (csgeNagII) for its. Our findings provide a new enzymatic tool and an effective strategy for releasing terminal β-1,2-GlcNAc at living cell level, which could facilitate the decoding of this glycan signal and support the modification of cell surface glycans.

This study opens a window for more studies. Besides emNagII, whether the other β-N-acetylhexosaminidases may also possess the potential to edit glycans on living cell surfaces merits further investigation. Moreover, emNagII also exhibits an activity in cleaving N-acetylgalactosamine in p-nitrophenyl β-D-GalNAc (data not shown), suggesting its utility for editing N-acetylgalactosamine modifications in living cell surface. It should be noted that the methodology employed in this study primarily targeted complex N-glycans across various substrates. As a portion of β-1,2-GlcNAc are masked beneath sialic acid and galactose, practical glycoengineering applications may require multi-enzyme collaboration, and further optimization of reaction conditions is required.

Furthermore, the complex biological functions of β-N-acetylhexosaminidases warrant further investigation. These enzymes enable microorganisms to degrade and utilize polysaccharides for growth and survival [50, 51]. Since β-N-acetylhexosaminidases was recognized as potential antifungal drug targets in 1997 [52], this concept has gained continued support. For instance, VhGlcNAcase is essential for *Vibrio marmoides* to degrade chitin and utilize GlcNAc as the sole carbon, nitrogen, and energy source. The growth of *V. marmoides* can be inhibited using N-acetylhexosaminidase inhibitors [53]. Similarly, BbHex1 from *Beauveria bassiana* enables the fungus to evade insect immune recognition by modifying the cell wall structure [54]. It also facilitates the degradation and penetration of the insect cuticle, thereby enhancing fungal pathogenicity. *E. meningoseptica* is a pathogen capable of infecting multiple species and causing serious life-threatening diseases in humans, such as meningitis, sepsis, and pneumonia [55, 56]. However, research on the pathogenic mechanisms underlying its infections remains limited. Based on these findings, we propose that emNagII may contribute to its survival and virulence. Further research on N-acetylhexosaminidases could provide new insights and strategies for understanding the pathogenic mechanisms of *E. meningoseptica* and other pathogens, as well as for developing potential therapeutic strategies.

## 5. Conclusion

In conclusion, we have identified a novel β-N-acetylglucosaminidase (emNagII) from *E. meningoseptica* and characterized its structural and enzymatic properties. Using emNagII, we accomplished the editing of terminal β-1,2-linked N-acetylglucosamine in complex N-glycans on living cell surfaces. This work provides the novel tool and insight to decode N-glycan signaling and enables direct glycan modification on living cell surfaces.

## Supporting information

Supplemental files will be used for the link to the file on the preprint site.

## Notes

### Competing Interest Statement

The authors have declared no competing interest.

### Funding

- National Key Research and Development Program of China (2021YFC 2700800) ; TMRC of Glycomics by Fudan Z & Sysdiagno (230016HZ003)

