## Supplemental files will be used for the link to the file on the preprint site. for "Identification and characterization of emNagII, a novel β-1,2-N-acetylglucosaminidase from *Elizabethkingia meningoseptica*"

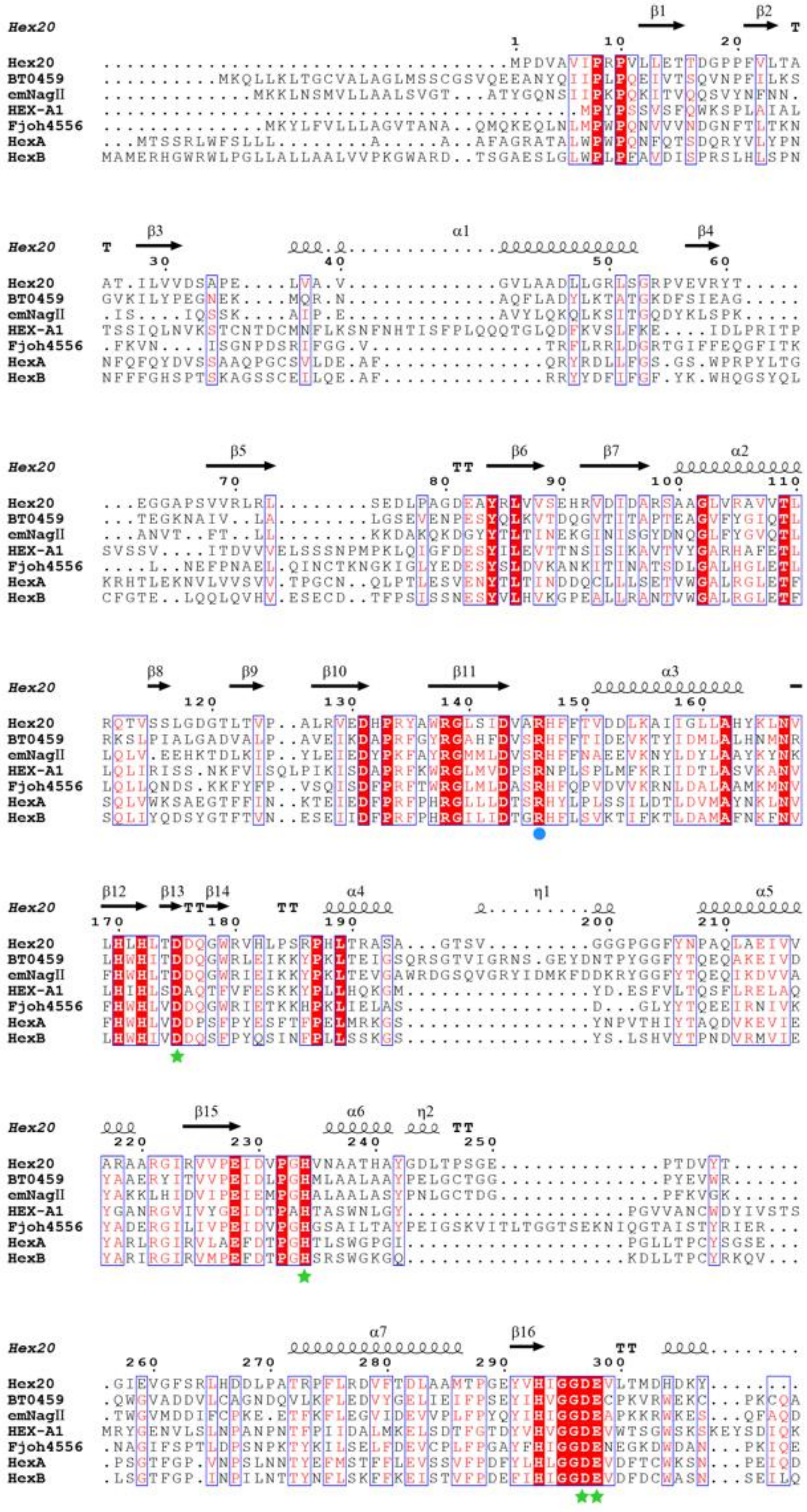
Hex20
BT0459
emNagII
HEX-A1
Fjoh4556
HexA
HexB

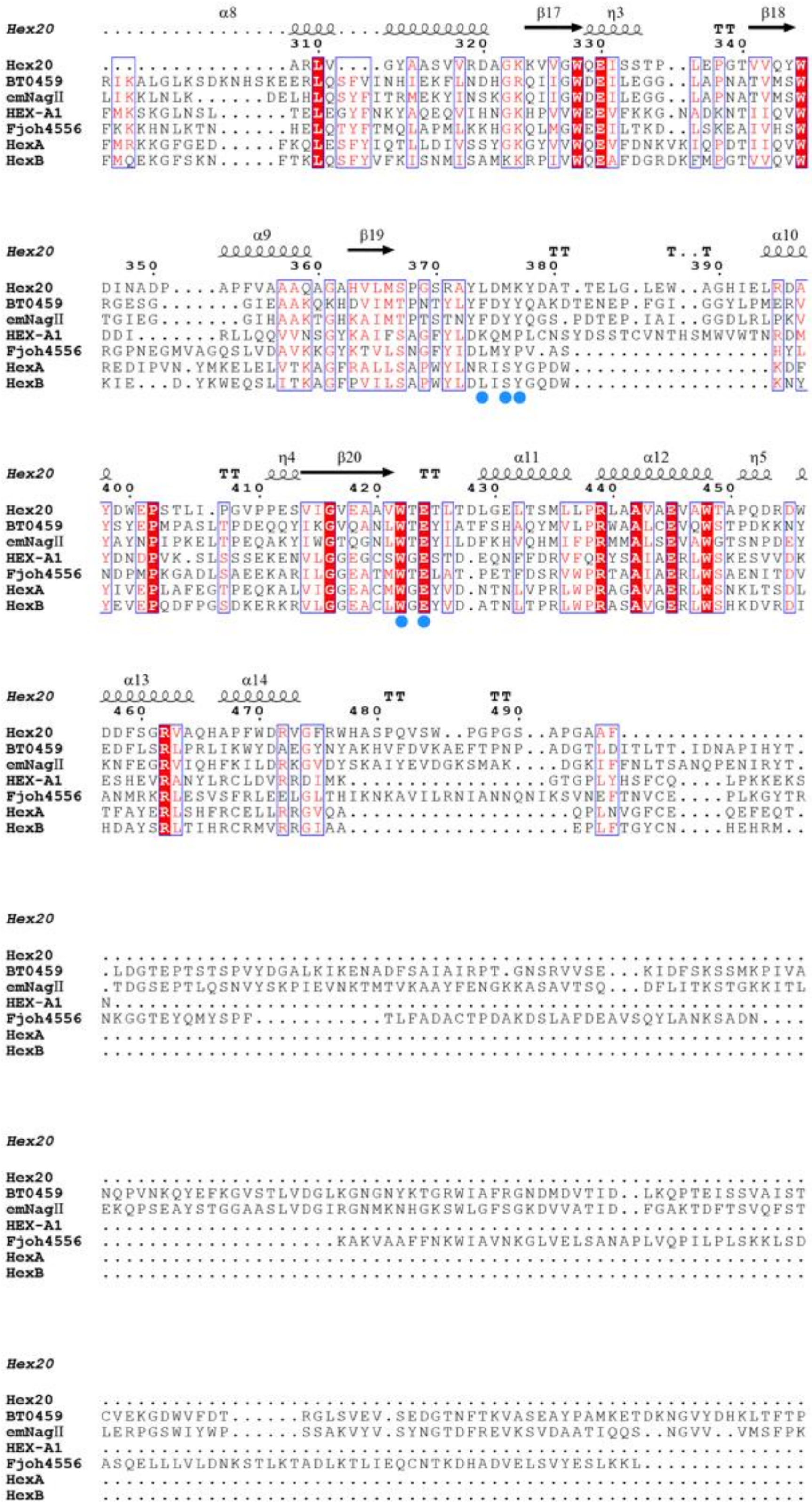
Hex20
BT0459
cmNagII
HEX-A1
Fjoh4556
HexA
HexB

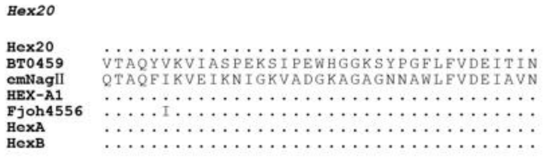


**Fig. S1 Sequence analysis of emNagII.** Structure-based sequence alignment of six GH20 family members using ESPript. Through multiple sequence alignment, the potential active sites were predicted and marked. Blue circles indicate potential binding sites, and green stars denote potential catalytic sites.

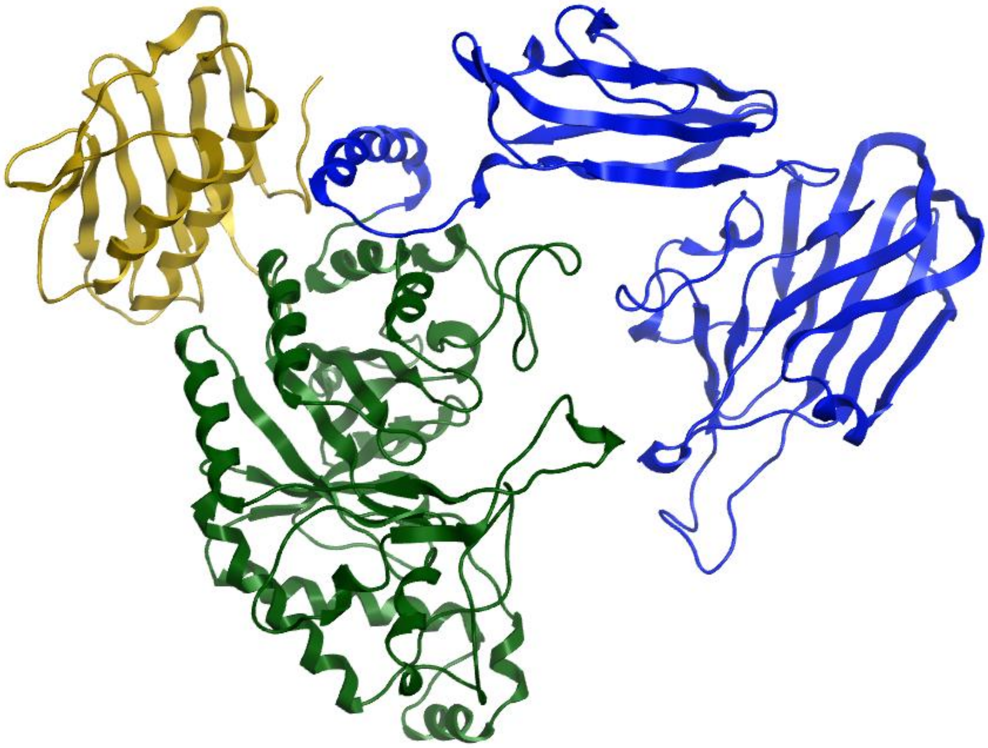

**Fig. S2 Predicted ribbon structure of emNagII by AlphaFold2.** The 727 amino acids of emNagII without signal peptide was input in to Alphafold2 and the structure was displayed by PyMOL. The N-terminal domain (1-122 aa) was colored in yellow, middle domain (123-467 aa) was in green and the C-terminal domain (468-727 aa) was in blue.
